# Assessment of photosynthetic activity in dense microalgae cultures using oxygen production

**DOI:** 10.1101/2023.11.29.569170

**Authors:** Antoni Mateu Vera-Vives, Tim Michelberger, Tomas Morosinotto, Giorgio Perin

## Abstract

Microalgae are photosynthetic microorganisms playing a pivotal role in primary production in aquatic ecosystems, sustaining the entry of carbon in the biosphere. Microalgae have also been recognized as sustainable source of biomass to complement crops. For this objective they are cultivated in photobioreactors or ponds at high cell density to maximize biomass productivity and lower the cost of downstream processes.

Photosynthesis depends on light availability, that is often not constant over time. In nature, sunlight fluctuates over diurnal cycles and weather conditions. In high-density microalgae cultures of photobioreactors outdoors, on top of natural variations, microalgae are subjected to further complexity in light exposure. Because of the high-density cells experience self-shading effects that heavily limit light availability in most of the mass culture volume. This limitation strongly affects biomass productivity of industrial microalgae cultivation plants with important implication on economic feasibility.

Understanding how photosynthesis responds to cell density is informative to assess functionality in the inhomogeneous light environment of industrial photobioreactors. In this work we exploited a high-sensitivity Clark electrode to measure microalgae photosynthesis and compare cultures with different densities, using *Nannochloropsis* as model organism. We observed that cell density has a substantial impact on photosynthetic activity, and demonstrated the reduction of the cell’s light-absorption capacity by genetic modification is a valuable strategy to increase photosynthetic functionality of dense microalgae cultures.

**Highlights:**

- Microalgae biomass is a promising alternative to crops.
- The impact of cultivation at scale on photosynthesis is still under-investigated.
- The Photosynthesis-Irradiance (PI) relationship is informative.
- High-sensitivity oxygen measurements for PI investigation was validated.
- The effect of cell density on PI was studied in *Nannochloropsis* pale mutants.

## 1. Introduction

Photosynthetic organisms are responsible for primary production, sustaining most lifeforms on our planet (Cavicchioli et al., 2019). Among them, eukaryotic microalgae are unicellular species that play a pivotal role in the biosphere, being responsible for approx. 50% of global primary production (Kirchman, 2018). Thousands of different microalgae species exist (Guiry, 2012), as the result of evolution in many ecological niches, where they are often placed at the basis of the existing food web, representing the primary producers and thus the main responsible of organication of atmospheric carbon.

Beside their fundamental ecological role, microalgae are also gaining increasing interest as bio-factories to convert the current fossil-fuels-based economy to a bio-based solar-driven alternative, for a more sustainable future (Olabi et al., 2023). Microalgae have a higher photosynthetic activity than plants, mainly because their whole organism is photosynthetically active. This leads to higher rates of CO_2_ and nutrients sequestration, enabling their cultivation in strict connection to industrial and civil sites to mitigate greenhouse gases emissions and water pollution (Ma et al., 2022; Molazadeh et al., 2019; Giorgio Perin et al., 2019). Moreover, their huge metabolic plasticity, developed in thousands of years of evolution in different ecological niches shaped the ability to accumulate a plethora of metabolites finding many applications in the current economy, e.g. from biofuels to food additives (Khan et al., 2018; Perin and Morosinotto, 2019a).

In microalgae, as in many other photosynthetic eukaryotes, the homeostasis of the central metabolism depends on the energetic and redox status of the cell, which is balanced by the activity of two organelles, i.e. chloroplasts and mitochondria. In strictly photoautotrophic microalgae species, mitochondrial respiration shows a minimal activity (10-15% of gross photosynthetic rate) and only sustains cell maintenance (Formighieri et al., 2012; Masojídek et al., 2021), making the photosynthetic activity of the chloroplast the final source of energy to sustain the central metabolism.

Chloroplasts contain membrane-bound enzymatic complexes that mediate photosynthesis, driving both i) the transfer of electrons from a reduced substrate (i.e. water) to an oxidized product (i.e. NADP^+^) and ii) the translocation of protons across a biological membrane to generate a proton motive force that fuels the synthesis of chemical energy in form of ATP by the action of ATP synthase.

Photosynthetic activity in the chloroplast depends on the availability of light energy, which in nature changes over diurnal cycles and weather conditions, affecting photosynthetic functionality and consequently microalgae fitness. The ability to respond to environmental changes in light availability is one of the phenomena at the base of the success of some microalgae species over others in different ecological niches, with implications for the functionality of the very ecosystem (Morgan-Kiss et al., 2006).

On the other hand, when microalgae biomass is to be exploited for industrial applications, these organisms are cultivated in open or closed systems, i.e. ponds or photobioreactors, respectively, that operate with high cell density to maximize biomass productivity (Ruiz et al., 2016). In such conditions, the high density leads to cell’s self-shading and prevents light to homogeneously reach all the regions of the mass culture, generating a light gradient from external to internal regions (Formighieri et al., 2012; Perin and Morosinotto, 2019b; Schediwy et al., 2019). The former receives excess light and the latter limiting irradiance, curbing global light-use efficiency in photobioreactors. Microalgae cultures at industrial scale are also actively mixed to i) increase the average exposition of the mass culture to light and ii) optimize nutrients and CO_2_ supply (de Souza Kirnev et al., 2022). Because of mixing, microalgae are constantly moved from most-exposed to light-limited regions of the culture (and *viceversa*) and experience fast fluctuations of irradiance to which regulatory mechanisms of photosynthesis are only partially able to respond to, affecting light-use efficiency and biomass productivity in photobioreactors (Bellan et al., 2020; Perin and Morosinotto, 2023). The impact of this complex light environment on microalgae physiology is still under-investigated (Masojídek et al., 2021), which is one of the most relevant causes for the limited success of the optimization efforts of microalgae cultivation at scale so far (Perin et al., 2022).

A more systematic investigation of how photosynthesis responds in high-density cultures, focusing e.g. on the influence of phenomena such as cell’s self-shading, is an unavoidable task to understand how microalgae deal with the complex environmental light conditions of photobioreactors and ultimately to deploy effective optimization efforts.

In this work, we used a high-sensitivity Clark electrode to measure the photosynthesis-irradiance relationship of microalgae cultures with different densities, using *Nannochloropsis gaditana* as experimental model. We quantified the impact of cell density on microalgae photosynthetic activity, showing how self-shading plays a relevant role in dense cultures. This hypothesis was validated by exploiting pale mutants accumulating less Chlorophyll per cell, which demonstrated the larger the reduction in pigment content, the higher the increase in photosynthetic activity.

## 2. Materials and methods

### 2.1 Microalgae strains and culture conditions

#### 2.1.1 Strains

In this work we used the microalgae species *Nannochloropsis gaditana,* strain CCAP 849/5, that was purchased from the Culture Collection of Algae and Protozoa (CCAP).

#### 2.1.2 Culture conditions

*N. gaditana* was maintained in F/2 solid media, containing 32 g l^-1^ sea salts (Sigma Aldrich), 40 mM Tris-HCl pH 8, Guillard’s (F/2) marine water enrichment solution (Sigma Aldrich) and 1% agar (Duchefa Biochemie).

Microalgae cells were pre-cultured in sterile F/2 liquid media in Erlenmeyer flasks exposed at 100 µmol of photons m^-2^ s^-1^ with 100 rpm agitation, at 22 ± 1 °C in a growth chamber. Growth curves were performed in the same conditions of pre-cultures from cells washed twice in fresh F/2 media, starting from 5·10^6^ cells ml^-1^ concentration in sterile F/2 liquid media supplemented with 10 mM NaHCO_3_ to avoid carbon limitation.

### 2.2 Preparation of cells for measurements of oxygen evolution

Cell concentration was measured with an automatic cell counter (Cellometer Auto X4, Cell Counter, Nexcelom) at the fourth day of cultivation in the growth conditions described above. Microalgae cells were collected via mild centrifugation at 3,500 g for 10 min at room temperature and then resuspended in fresh sterile F/2 media supplemented with 10 mM NaHCO_3_ right before the start of oxygen evolution assessment to avoid carbon limitation during the measurement.

### 2.3 High resolution oxygen evolution

Oxygen evolution was measured at the fourth day of the growth curves described above. Measurements were performed using a test version of the NextGen-O2k and the PhotoBiology (PB)-Module (Oroboros Instruments, Innsbruck) with the software DatLab 7.4.0.4 (Went et al., 2021), according to the methods developed in (Vera-Vives et al., 2022). The PB light source contained a blue OSLON® LED (emitting wavelength range 439-457 nm with the peak at 451 nm, manufactured by OSRAM) attached to the window of the NextGen-O2k chamber.

The oxygen concentration was assessed in 2-ml measuring chambers at 22 °C with a 2-seconds frequency and samples were magnetically stirred at 750 rpm.

Two measurements were performed in parallel at each time, taking advantage of the two chambers of the instrument.

At first, the measuring chambers were filled with fresh and sterile F/2 medium, containing 10 mM NaHCO_3_ to avoid carbon limitation during the measurement and to equilibrate the temperature at 22 °C for few minutes. Then, a fraction of the medium (<10% of the chamber volume) was replaced with a suspension of *Nannochloropsis* cells to reach the desired final concentration in the measuring chamber. The chambers were then closed, and the samples were dark adapted (at least for 10 min until the oxygen consumption rate was constant) to assess the respiration rate before starting the measurements of the oxygen flux at increasing irradiances.

After stabilization of the respiration signal, light was turned on at increasing irradiances, waiting at least 5 min at each light intensity to achieve the stabilization of the oxygen evolution rate (on average the O_2_ flux trace stabilized within the first 2-3 min, Supplementary Figure S1). The values of oxygen evolution rate reported in this work at each irradiance correspond to the median of 40-50 points in the stable region of oxygen flux.

Respiration and photosynthesis rates at different irradiances were calculated with the software DatLab 7.4.0.4. It is worth noting that the photosynthetic parameters extrapolated from the photosynthesis-irradiance curves were not influenced by the starting concentration of molecular oxygen, as a preliminary equilibration phase of the measuring medium in presence of atmospheric oxygen was carried out for all tested samples.

### 2.4 Determination of chlorophyll content

After evaluation of oxygen evolution, microalgae samples were further processed to extract chlorophyll molecules from *Nannochloropsis* cells, using 1:1 ratio of 100% N, N-dimethylformamide (DMF) (Sigma Aldrich), at 4 °C in the dark, for at least 24 h (Wellburn, 1994). The chlorophyll concentration was calculated, using specific extinction coefficients (Wellburn, 1994), from the absorption values at 664 nm of DMF *Nannochloropsis* extracts, collected using a Cary 100 spectrophotometer (Agilent Technologies).

### 2.5 Statistical analysis

In this work, we performed a statistical hypothesis testing for all the data presented. Statistical significance was assessed by t-test using OriginPro 2020 (v. 9.7.0.188) (http://www.originlab.com/). Samples size was at least 4 for all the measurements collected in this work. Data were fitted with the equation defined in (Ye, 2007), using a minimum mean square error-based approach using OriginPro 2020 (v. 9.7.0.188).

## 3. Results

### 3.1 Estimation of photosynthetic functionality from oxygen evolution

*Nannochloropsis* cells grown in lab-scale cultures, as described in the materials and methods section, were used to investigate the photosynthesis-irradiance (PI) relationship of samples with increasing cell concentration. Figure 1 shows data obtained with 5 · 10^6^ cells ml^-1^, corresponding to a chlorophyll concentration of 4 μg Chl ml^-1^. Lower concentrations (down to 1 · 10^6^ cells ml^-1^) have a higher noise but still were able to generate traces of sufficient quality for a reliable extrapolation of the parameters describing the photosynthetic activity of *Nannochloropsis* cells (Supplementary Figure S2).

**Figure 1.**
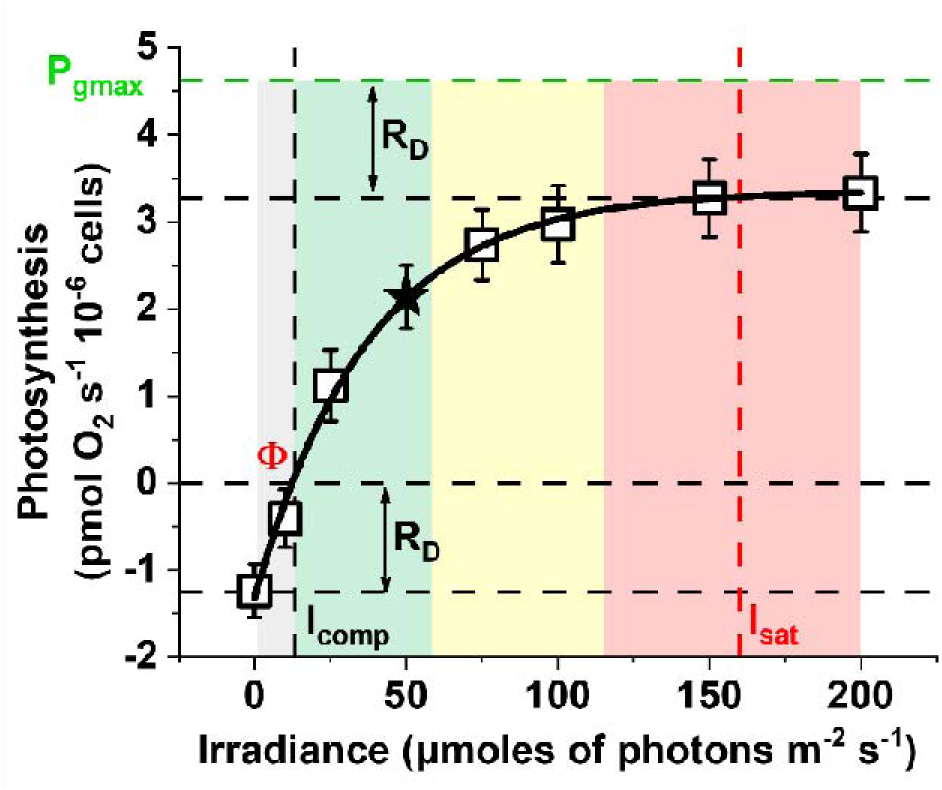
Dependence of Photosynthesis on light intensity. Data shown here were assessed from a *Nannochloropsis* culture with 5 · 10^6^ cells ml^-1^ cells concentration, corresponding to 4 μg Chl ml^-1^. Photosynthesis is expressed as O_2_ flux and it is normalized to the number of cells. Light intensity is expressed as irradiance. and indicate the light compensation and light saturation points, respectively, both expressed as [μmol photons·s^-1^·m^-2^]. is the dark O_2_ respiration rate, expressed as [pmol·s^-1^·10^-6^ cells], is the maximal gross O_2_ photosynthetic rate, expressed as [pmol·s^-1^·10^-6^ cells]. is the quantum yield in the range between and [pmol O_2_ ·m^2^·μmol photons^-1^·10^-6^ cells]. The black star indicates the photosynthetic activity (expressed as O_2_ flux) at 50 μmol photons s^-1^ m^-2^, a parameter used in the following analysis. In this scheme, four irradiance regions have been highlighted with different colours: grey, the respiration rate is higher than the photosynthetic rate; green, the photosynthetic rate linearly increases with irradiance; yellow, the photosynthetic rate increases with irradiance up to a saturation limit; red, the photosynthetic rate does not increase with irradiance. Every point in this figure represents the average value of 4 biological replicates ± SD. The data for each biological replicate at each irradiance come from the median value of 40 datapoints where the O_2_ trace was more stable (Supplementary Figure S1).

*Nannochloropsis* samples at a concentration of 5 · 10^6^ cells ml^-1^ showed a dark respiration rate (of 1.31 ± 0.12 pmol O_2_ s^-1^·10^-6^ cells. Light intensity was then progressively increased from 0 to 200 μmol photons s^-1^ m^-2^. The light compensation point (i.e. I_comp_, namely the irradiance value where the dark respiration is fully compensated by the photosynthetic activity rate) was 12.2 ± 1.1 μmol photons s^-1^ m^-2^, while the saturation threshold (i.e. I_sat_, namely the irradiance value at which the photosynthetic rate does not increase with irradiance any longer) was 163 ± 27 μmol photons·s^-1^·m^-2^. The maximal gross photosynthetic rate was instead 4.6 ± 0.2 pmol O_2_ s^-1^·10^-6^ cells.

In this work we exploited a high-sensitivity Clark electrode (i.e. NextGen-O2k) for quantifying photosynthesis that enables higher resolution compared to alternative versions. The NextGen-O2k has a resolution of 2 nM and the limit of detection of the oxygen flux is 0.001 µM s^-1^ (Doerrier et al., 2018; Gnaiger, 2008), therefore enabling the reliable detection of small differences in O_2_ concentration. It is in fact worth noting that we could decrease the cell concentration by a 30x factor, compared to similar experiments with an alternative Clark-type sensor (Perin et al., 2015). Consequently, we could determine the photosynthetic parameters above detailed with relatively low-concentration microalgae samples, without sacrificing the reproducibility. The minimization of the volume of the microalgae culture has a twofold important implication: i) self-shading effects during the measurements are minimized for an unbiased determination of photosynthetic parameters and ii) changes in oxygen flow per volume of culture are relatively small, enabling to run even longer-time experiments without reaching saturation with possible negative effects on microalgae cells.

### 3.2 Dependance of photosynthetic functionality on cell concentration

Thanks to the instrumentation sensitivity it was even possible to assess the impact of self-shading by measuring samples of *Nannochloropsis* with increasing cell concentrations, from 2.5 to 100 · 10^6^ cells ml^-1^ (Figure 2). For a fair comparison, we used the photosynthetic activity data collected at 50 µmol photons m^-2^ s^-1^, namely an irradiance value within the linear correlation range between the O_2_ flux and light intensity in the PI curve of Figure 1 (black star in Figure 1).

**Figure 2.**
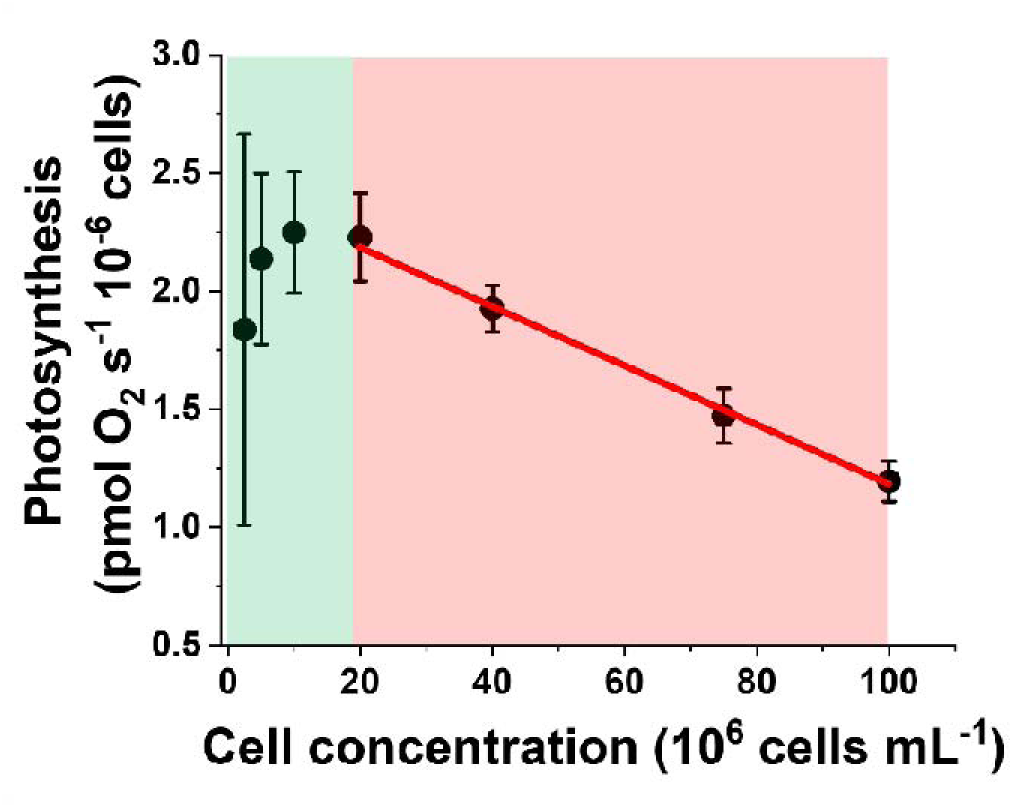
Photosynthetic activity as a function of cell concentration. Photosynthesis is expressed as net O_2_ flux and it is normalized to the number of cells. The net O_2_ flux here reported corresponds to the values measured at 50 µmol photons m^-2^ s^-1^ (i.e. the linear phase of correlation between photosynthesis and irradiance in the PI curves of Supplementary Figure S2) for *Nannochloropsis* samples with increasing cell concentration: 2.5, 5, 10, 20, 40, 75 and 100 · 10^6^ cells ml^-1^, and corresponds to the star value in Figure 1. The O_2_ flux is constant in the range 2.5 – 20 · 10^6^ cells ml^-1^ (green area), whilst it decreases as the cell concentration increases between 20 – 100 · 10^6^ cells ml^-1^ (red area). The slope of the linear correlation equation in the green area is not significantly different from 0 [y = (2.12 ± 0.11) + (0.006 ± 0.007) x, Pearson’s R: 0.5, R^2^: 0.25], whilst in the red area it is [y = (2.43 ± 0.03) – (0.01 ± 0.0004) x, Pearson’s R: -0.99, R^2^: 0.99 (t-Test, p-value < 0.05). Data refer to the average ± SD of four independent biological replicates.

All cells were exposed to the same light conditions during cultivation ensuring an identical acclimation state (average Chl content of 0.08 µg Chl · 10^-6^ cells).

From 2.5 to 20 · 10^6^ cells ml^-1^ samples showed a constant O_2_ flux, whilst the latter decreased as the cell concentration increased up to the highest concentration tested in this work (100 · 10^6^ cells ml^-1^, Figure 2). Overall, cell concentrations could be divided into two groups according to the behavior observed in Figure 2: i) for samples between 2.5 and 20 · 10^6^ cells ml^-1^ there was no significant difference between the measured photosynthetic activity, suggesting that in this range data are not influenced by the cell concentration and thus the self-shading effects during the measurements are neglectable. In the former case, with 2.5 ·10^6^ cells ml^-1^, the noise of the measurements was higher because of the low cell concentration. ii) for samples in the range between 40 and 100 · 10^6^ cells ml^-1^, photosynthetic activity instead decreased as the cell concentration increased. This is likely to depend on the inhomogeneous light distribution in the samples because of their high optical density. Maximal gross photosynthesis values are equal for all concentrations measured, as expected since all cells are saturated. This confirms that the differences of photosynthetic activity observed at lower intensities depend on a shading effect (Supplementary Table S1).

### 3.3 Measuring photosynthetic functionality in microalgae dense cultures

To better assess how self-shading affected photosynthetic activity of *Nannochloropsis* cells, the photosynthesis-irradiance relationship of diluted and dense samples (5 and 100 · 10^6^ cells ml^-1^, respectively) were compared (Figure 3).

**Figure 3.**
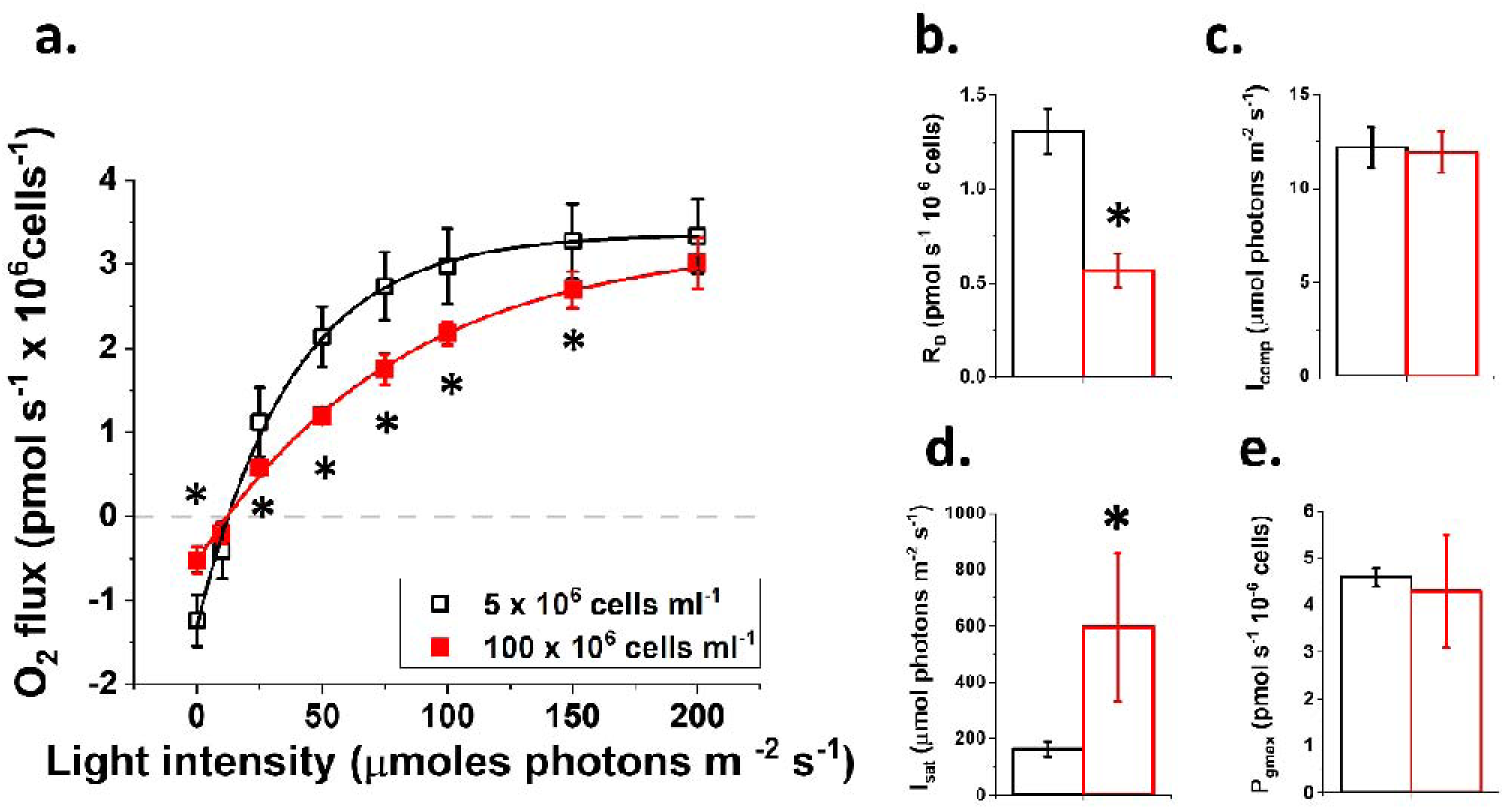
Photosynthesis-Irradiance dependance as a function of culture density. The photosynthesis-Irradiance relationship was measured for diluted and dense *Nannochloropsis* cultures (i.e. 5 and 100 · 10^6^ cells ml^-1^, respectively) and fitted with the equation defined in (Ye, 2007), using a minimum mean square error-based approach (a). The fitting returned the mathematical parameters indicated in panel from b) to e), that describe the differences in the shapes of the curves represented in panel a). For each mathematical parameter, the asterisk indicates a statistically significant difference between the two culture densities (t-Test, p-value < 0.05). Data refer to the average ± SD of four independent biological replicates. Black, 5 · 10^6^ cells ml^-1^ and red, 100 · 10^6^ cells ml^-1^ cell concentration. The same parameters for all the other cell concentrations tested in this work are reported in Supplementary Table S1.

We observed that the concentration of cells had a substantial effect on the shape of the photosynthesis-irradiance relationship, especially during the linear phase (Figure 3a). The dark respiration rate (R_D_) of the densest sample (100 · 10^6^ cells ml^-1^) was <30% of the value measured for the diluted sample (5 · 10^6^ cells ml^-1^, Figure 3b), whilst no significant differences were observed for the light compensation point (I_comp_) (Figure 3c). The lower RD might depend on a lower diffusion efficiency of O_2_ in the sample at high cell density. The light saturation point (I_sat_) was instead higher in the densest sample (Figure 3d). This can be explained because of the higher optical density and cells shading in the sample, that thus requires stronger illumination to reach saturation. This hypothesis was confirmed by the observation that the maximal gross O_2_ photosynthetic rate (P_gmax_) was the same for both cell concentrations (Figure 3e). The value of maximal gross photosynthesis is therefore reliably estimated even in dense microalgae samples, because when light is in excess the effect of self-shading becomes negligible since all cells are exposed to saturating illumination.

Overall, the methodology employed in this work enables the reliable estimation of photosynthetic parameters both in low and high optical density microalgae samples. In the second case, this enables to extrapolate the photosynthetic performances of microalgae cultures, like those typical of industrially relevant cultivation plants, already at the lab-scale.

### 3.4 How does cell’s light absorption capacity impact the Photosynthesis-Irradiance relationship in microalgae dense cultures?

The methodology described in this work offers the opportunity to investigate the photosynthetic activity of microalgae in complex light environments and quantitatively assess the impact of shading on photosynthetic activity. In the past few years, we successfully isolated *Nannochloropsis* mutants with different degrees of reduction in the Chl content [Figure 4a, (Perin et al., 2015)], which did not result in any affected phototrophic growth phenotype at the lab-scale (Figure 4b). These strains indeed represent a useful tool to better describe how microalgae photosynthesis responds when shading effects, typical of cultivation in high optical-density conditions, become relevant.

**Figure 4.**
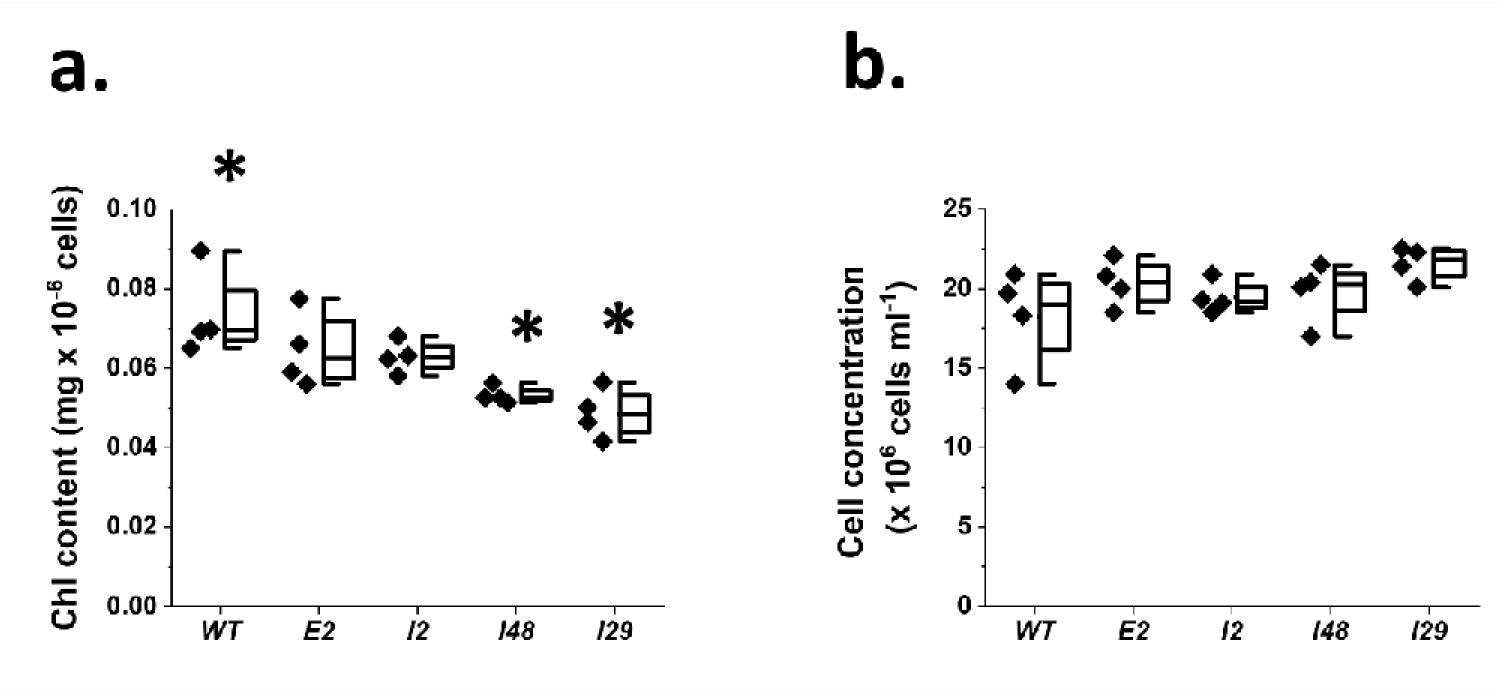
*Nannochloropsis* mutants with a reduced chlorophyll content and unaffected phototrophic growth. *Nannochloropsis* mutants showing different degrees of reduction of chlorophyll (Chl) content (a), but an affected phototrophic growth (b) with respect to the parental strain (WT) were isolated (Perin et al., 2015). Data refer to the average ± SD of four independent biological replicates, grown in plastic cell culture flasks with ventilation for four days, according to the protocol detailed in the materials and methods section. Asterisks indicate statistically significant differences between one mutant and the parental strain (Test-t, p-value < 0.05).

In this work, we measured the photosynthesis-irradiance relationship of these microalgae strains and compared it to the reference parental *Nannochloropsis* strain, in both diluted and dense cultures (Figure 5a and 5b). In this case, to account for the different Chl content of the strains under investigation, data were normalized on the Chl content (Figure 5). We observed that the effect of cell concentration on the shape of the photosynthesis-irradiance relationship, that we previously measured for the WT (Figure 3a), is maintained also in photosynthetic pale mutants (Supplementary Table S3).

**Figure 5.**
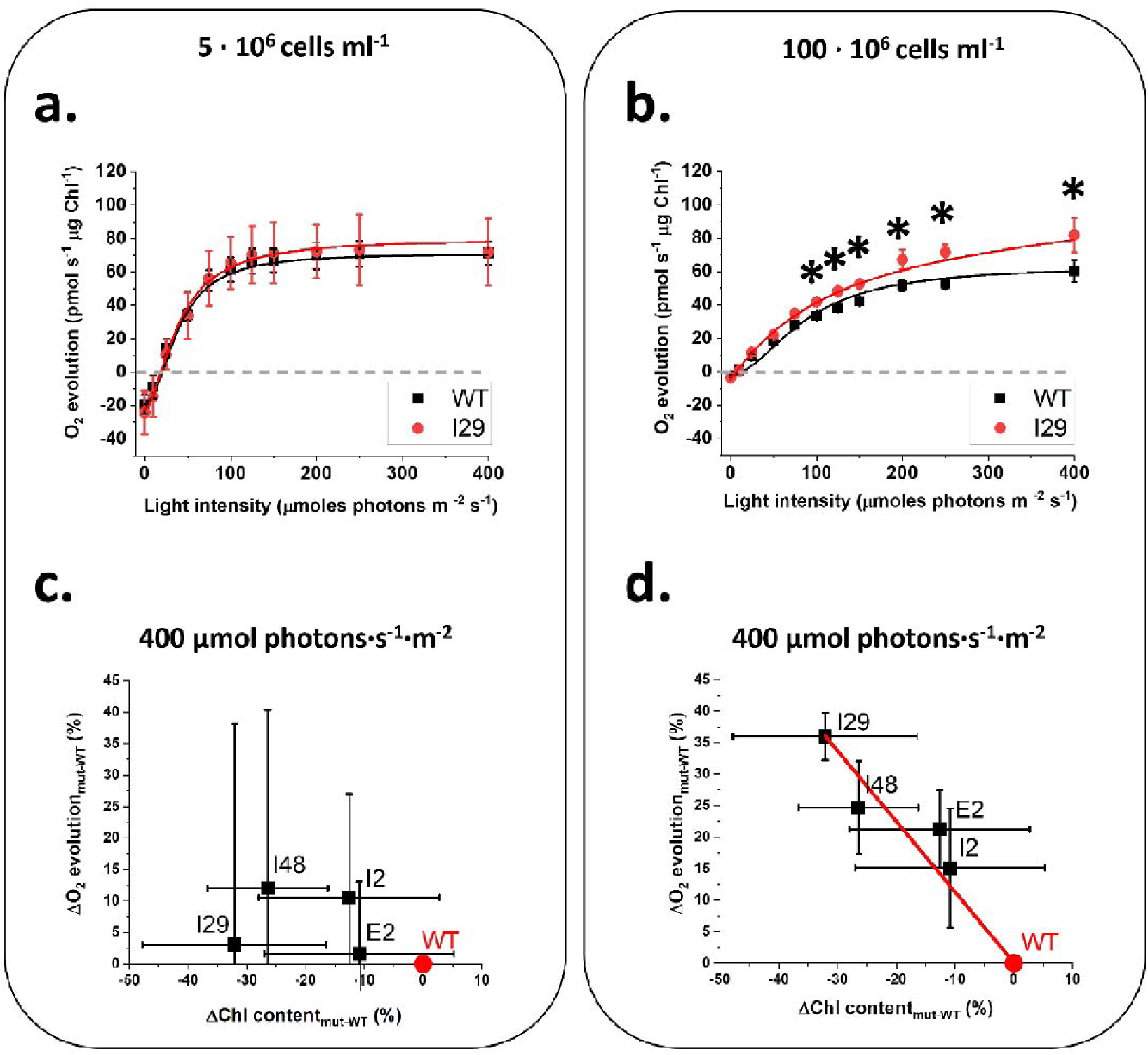
Improvement of photosynthetic activity as a function of the reduction of the pigment content, in *Nannochloropsis* cultures with different densities. Examples of Photosynthesis-Irradiance relationship measured for *Nannochloropsis* WT and I29 strain, showing 30% reduction of Chl content with respect to the former (Fig. 4a), in diluted (a) (i.e. 5 · 10^6^ cells ml^-1^) and dense (b) (i.e. 100 · 10^6^ cells ml^-1^) cultures. The Chl content of the strains used in this experiment is reported in Figure 4a. Data were fitted with the equation of (Ye, 2007) using a minimum mean square error-based approach. Improvement in O_2_ flux at 400 μmol photons s^-1^ m^-2^, expressed as percentage with respect to the value of the parental strain (WT) for the four mutants showing different degrees of reduction in the chlorophyll (Chl) content, expressed as percentage of reduction with respect to the WT, for diluted (c) and dense (d) *Nannochloropsis* cultures. Data correspond to the photosynthetic activity measured at 400 μmol photons·s^-1^·m^-2^, that is the irradiance to which dense cultures in lab-scale PBRs demonstrated an improved biomass productivity (Perin et al., 2017a). There isn’t linear correlation between x and y observations in panel c), whilst there is linear correlation in panel d) (red continuous line), (t-Test, p-value < 0.05). Parametric values for the correlation equations for the two panels are reported in Supplementary Table S2. Data refer to the average ± SD of four independent biological replicates. Asterisks indicate statistically significant differences between one mutant and the parental strain (t-Test, p-value < 0.05). The original data used in this figure are reported in Supplementary Figure S3.

Also in this case, the dark respiration rate (RD) of dense cultures was lower than in diluted samples, with implications also on the light saturation point. Densest cultures showed a higher light saturation point than diluted samples, consistently with shading (Figure 5a, 5b and Supplementary Table S3). Overall, these data suggest the photosynthetic mutant strains investigated in this work respond to the environmental conditions of dense cultures, as well as the parental strain.

In both diluted and dense cultures, the mutant strains here investigated showed an increase in the dark respiration rate (R_D_) with respect to the WT, as expected (Formighieri et al., 2012). No major differences in other photosynthetic parameters were instead observed between WT and mutant strains in diluted conditions, whilst in dense cultures we observed a shared increase in P_gmax_, I_comp_ and I_sat_ (Supplementary Table S3), as expected on a Chl basis for Chl-less strains (Formighieri et al., 2012).

Strain I29, that, out of the four strains tested in this work, had the largest reduction in pigment content with respect to the WT, did not show any difference of photosynthetic functionality than the WT in diluted cultures (Figure 5a). In dense cultures, I29 instead showed with equal cell concentration a higher photosynthetic activity with respect to the WT at most of the tested irradiances, with the difference increasing with light intensity. In these conditions, I29 also saturates photosynthesis at higher irradiances than the parental strain (Figure 5b). These data suggest that mutants with a reduced pigment content show an improvement of photosynthetic functionality vs. the parental strain, but it depends on the density of the culture, where self-shading effects likely become relevant for photosynthesis. To validate this hypothesis, we measured the Photosynthesis-Irradiance relationship of the other three pale mutants presented in Figure 4. It is in fact worth noting that the four pale *Nannochloropsis* mutants of this work showed a progressively larger reduction of pigment content than the WT (Figure 4a), making them the ideal choice to assess the impact of a wide range of shading degrees because of differences in the cells’ light absorption capacity. We observed that also the other three strains tested in this work showed an improved photosynthetic functionality with respect to the WT and that this phenomenon is observable only in dense cultures, but in these cases only at high irradiance (i.e. 400 μmol photons·s^-1^·m^-^2, Supplementary Figure S3). It is worth remembering that these strains showed a lower reduction of Chl content than the WT, with respect to strain I29 (Figure 4a). It is therefore likely that a reduced cell’s light absorption capacity can drive an improved light distribution profile in a dense culture, but the latter phenomenon is understandably influenced by the presence of a relevant shading effect, which in turn depends not only on the culture density alone, but also on the incident light.

To validate the latter conclusion, we plotted the improvement of photosynthetic activity measured at 50 μmol photons·s^-1^·m^-2^ (i.e. expressed as percentage with respect to the photosynthetic activity of the WT), for the four pale mutants investigated this work, versus the reduction of pigment content, as percentage of the pigment content of the WT (Supplementary Figure S4) and compared diluted and dense cultures. 50 μmol photons·s^-1^·m^-2^ was chosen for an unbiased comparison between strains because it falls in the linearity range of correlation between photosynthetic activity and irradiance (Figure 1).

We observed that at this irradiance there wasn’t any improvement of photosynthetic activity for any of the four strains here investigated in diluted conditions with respect to the WT (Supplementary Figure S4a). Strain I29 was the only one to show an improved photosynthetic activity in dense conditions, but overall, there was no relevant linear correlation between improvement in photosynthetic activity and reduction of pigment content either (supplementary Figure S4b).

When we plotted the improvement of photosynthetic activity measured at 400 μmol photons·s^-1^·m^-2^, versus the reduction of pigment content for the four pale mutants investigated this work, instead we observed a significant difference between diluted and dense conditions. In diluted cultures there was no improvement in photosynthetic activity (Figure 5c), whilst in dense conditions, there was a significant correlation between improvement in photosynthetic functionality and reduction in pigment content, with the greatest reduction of the latter leading to the greatest improvement of photosynthetic activity (Figure 5d).

It is worth mentioning that the strains investigated in this work showed an improved biomass productivity in dense lab-scale PBR cultures exposed at 400 μmol photons·s^-1^·m^-2^ indeed, with the greatest advantage observed for the strains showing the strongest reduction in the Chl content (i.e. strain I29) (Perin et al., 2017a).

Overall, these data demonstrate that photosynthetic activity is affected by the cultivation conditions, with shading effects of dense cultures making a substantial difference with respect to diluted conditions. Shading effects can be effectively reduced by decreasing cell’s light absorption capacity, leading to a significant improvement in light distribution and consequently photosynthetic activity, provided enough photons are used to drive an inhomogeneous light distribution profile in dense WT microalgae cultures.

## 4. Discussion

### 4.1 Estimation of PI relationship in both diluted and dense microalgae cultures

Historically, the functionality of the photosynthetic metabolism has been investigated using the rate of carbon fixation in ^14^C-treated samples (Jassby and Platt, 1976; Perry et al., 1981). This methodology is complicated by the need of marked carbon which is expensive and requires a complex experimental set-up for the homogenous treatment of the sample under investigation. The development of CO_2_ detectors opened the possibility to measure the rate of carbon fixation avoiding the use of marked carbon. Nevertheless, this methodology is still limited by operational constraints like i) the sensitivity of the detectors that is currently too low to obtain reliable information in carbon limiting conditions and ii) the need of complex experimental set-ups (e.g. closed chambers) that can affect versatility of the method.

Molecular oxygen is the other product of photosynthesis, and its concentration is directly proportional to the number of electrons that are vehiculated through the photosynthetic electron transport chain, making the rate of O_2_ evolution a good choice for the investigation of photosynthetic functionality.

The quantification of the concentration of molecular oxygen has been originally achieved with the so-called clear and dark bottle method (Strickland, 1960). Afterwards, devices based on Clark-type electrodes have been developed, yet the information achievable have been limited by the i) low resolution and sensitivity and ii) difficulty in the precise control of light supply to the sample.

In this work we used one high-sensitivity Clark electrode to improve both these constraints and demonstrated that it can detect the concentration of molecular oxygen in very-low-concentration samples (e.g. 1 and 2.5 · 10^6^ cells ml^-1^, Supplementary Figure S2) in *Nannochloropsis*. These concentration values are on average 30 times lower than those used with alternative Clark sensors (Ben-Sheleg et al., 2021; Perin et al., 2015; Vonshak et al., 2020), minimizing the volume of sample to sacrifice for the evaluation of photosynthetic activity of microalgae cultures. More importantly, the investigation of photosynthetic functionality at low cell concentration opens the possibility to investigate cultures with minimal shading.

Indeed, we were able to assess the impact of different cell densities on microalgae photosynthetic functionality. When we compared the Photosynthesis-Irradiance (PI) relationship of *Nannochloropsis* cultures at different concentrations, from diluted (i.e. 5 · 10^6^ cells ml^-1^) to dense (100 · 10^6^ cells ml^-1^), we observed that there was an effect of cell density on the shape of the PI curve (Figure 3a). At non-saturating irradiances [<150 μmol photons·s^-1^·m^-2^ for *Nannochloropsis*, (Sforza et al., 2012)], the photosynthetic activity was lower in dense cultures, indeed suggesting the high cell concentration triggers an inhomogeneous light distribution profile within the sample, with a substantial fraction of cells exposed to limiting light and thus not receiving enough energy to drive photochemistry. Nevertheless, maximal gross photosynthesis in the high-density sample equals the diluted culture, indicating the method could still reliably estimate photosynthetic parameters even in dense microalgae cultures (Figure 3e), enabling extrapolation of photosynthetic performances in cultivation conditions typical of industrial systems already at the lab-scale.

### 4.2 Cells’ light absorption capacity affects photosynthetic functionality in dense microalgae cultures

The method employed in this work enables to quantitatively assess the impact of cell’s self-shading on microalgae photosynthetic activity. To investigate this phenomenon in detail, we measured the PI relationship in microalgae strains with a significant reduction in their Chl content with respect to the parental strain. We used four *Nannochloropsis* strains, previously isolated for a reduced Chl content and unaffected phototrophic growth (Figure 4) and measured their PI curves using one high-sensitivity Clark electrode, comparing diluted and dense cultures (Figure 5). We observed that all mutant strains responded to the culture density as well as the WT and showed a reduction of the dark respiration rate (R_D_) and an increase of I_sat_ as the density increased (Supplementary Table S3). We hypothesize the lower R_D_ might depend on a lower diffusion efficiency of O_2_ in the sample at high cell density, likely driving to transient hypoxia regions in the mass culture volume.

In both diluted and dense conditions, the mutant strains showed an increased dark respiration rate (R_D_), I_comp_, I_sat_ and P_gmax_ with respect to the WT, as expected on a Chl basis from prediction models for Chl-less mutants (Formighieri et al., 2012).

At limiting light (i.e. 50 μmol photons·s^-1^·m^-2^), we didn’t observe any improvement of photosynthetic activity either in diluted or dense cultures (Supplementary Figure S4), suggesting two hypothesis: i) in diluted cultures, cell concentration was not high enough to drive a different light profile between mutants and WT cultures (Figure 6a), whilst ii) in dense conditions, the number of photons was not enough to drive a significant difference in the light distribution profile between mutants and WT cultures (Figure 6b).

**Figure 6.**
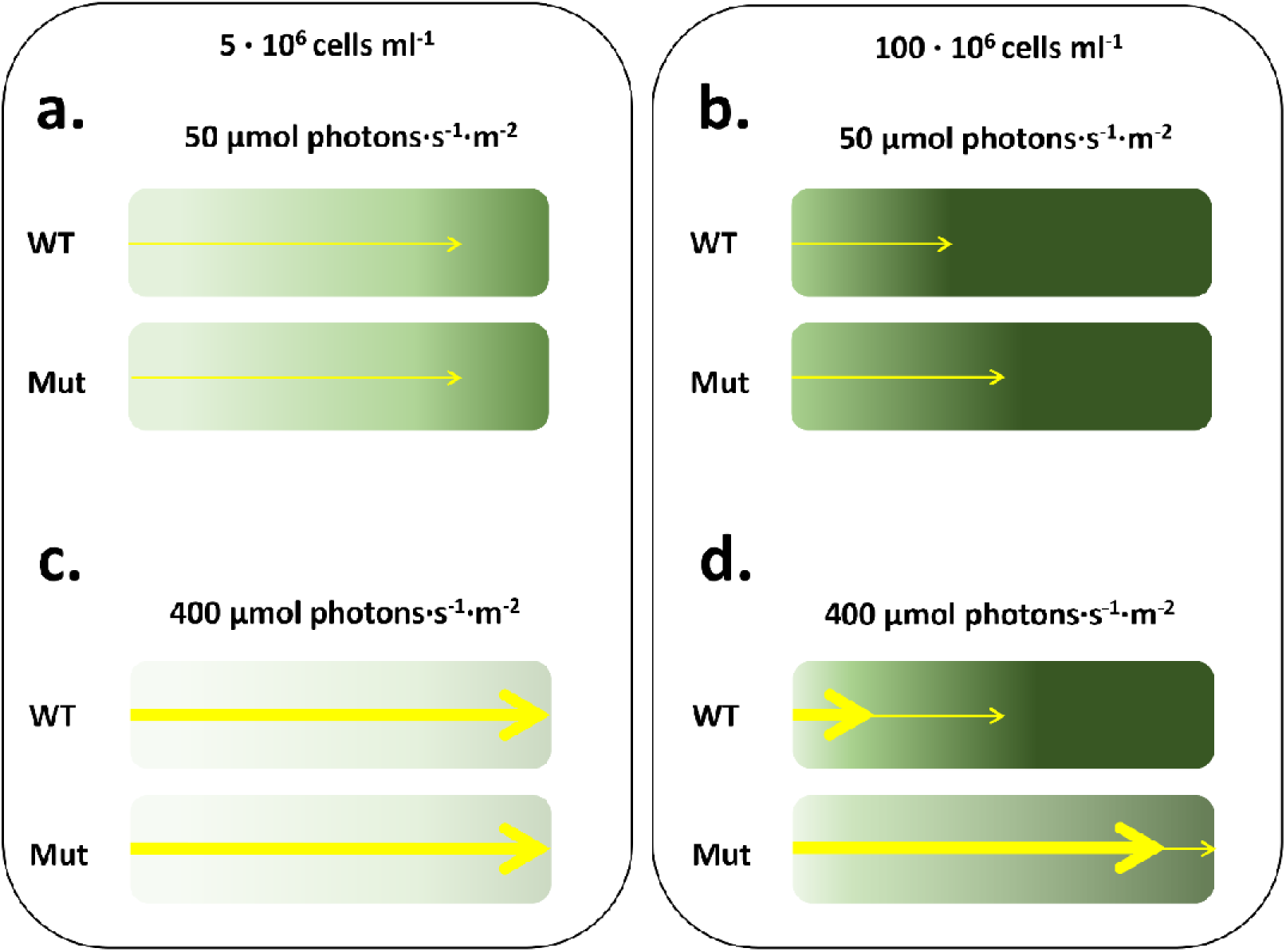
Light distribution profile in microalgae cultures. Schematic representations of the light distribution profile (yellow arrows) in WT and pale green mutants (Mut) *Nannochloropsis* cultures, in diluted (a and c) and dense (b and d) cultures, exposed at limiting (i.e. 50 μmol photons·s^-1^·m^-2^, a and b) and saturating (i.e. 400 μmol photons·s^-1^·m^-2^, c and d) light intensity.

At saturating light (i.e. 400 μmol photons·s^-1^·m^-2^), we still did not observe any significant difference in photosynthetic activity vs the WT for any of the four strains here investigated in diluted samples (Figure 5c), whilst in dense cultures we observed a significant improvement in the O_2_ flux with respect to the parental strain, that progressively increased with the reduction of the Chl content (Figure 5d). 400 μmol photons·s^-1^·m^-2^ is the irradiance at which the four mutant strains indeed showed an improved biomass productivity than the WT in the high density condition tested in this work (Perin et al., 2017a). This is likely to depend on a more homogeneous light distribution in dense cultures of microalgae with a reduced Chl content because of a reduced self-shading, with more light reaching the cells most distal from the light source than in WT cultures (Formighieri et al., 2012; G. Perin et al., 2019) (Figure 6d).

This phenomenon is expected to drive a higher photochemical rate in these regions of the culture volume, with beneficial consequences on the growth of the whole mass culture. Such improvement is observable only in dense microalgae cultures, because in diluted conditions light is likely to be homogeneously distributed even for WT cells (Figure 6c).

Along the rationale that a reduced pigment content can improve the light distribution profile within a mass culture, in the past years, several research efforts were successfully devoted to the isolation of paler microalgae strains, in different species. In controlled lab-scale conditions, these strains were demonstrated to increase light-use efficiency and growth with respect to the reference parental strains indeed (Cazzaniga et al., 2014; Dall’Osto et al., 2019; Kirst et al., 2012; Kirst and Melis, 2014; Mussgnug et al., 2005; Perin et al., 2015). Also the four mutant strains used in this work were demonstrated to have an improved biomass productivity in lab-scale PBRs (Perin et al., 2017a).

However, once pale mutant strains were tested at a higher cultivation scale (e.g. pilot plants outdoors), mixed results were obtained, with the confirmation of a growth advantage in some cases (Cazzaniga et al., 2014), but not in others (De Mooij et al., 2014). This phenomenon is yet to be fully addressed but is shedding light on the role played by the cultivation environment on photosynthesis. As observed e.g. for domesticated *Nannochloropsis* strains (Perin et al., 2017b), photosynthetic phenotypes acclimate to the cultivation environment and they can respond with a different intensity than the corresponding parental strain. Consequently, the cultivation environment, in some cases, can even mask the photosynthetic features isolated at the lab-scale and the original advantage can be lost. This discrepancy of behaviors from the lab- to the industrial-scale of cultivation has been under-investigated so far, mainly because most of the phenomena impacting microalgae photosynthesis in photobioreactors can only be assessed at scale (Masojídek et al., 2021), calling for, often unbearable, economic efforts.

The methodology exploited in this work opens the possibility to extrapolate the photosynthetic functionality of microalgae in dense cultures typical of industrial cultivation plants already at the lab-scale, enabling the collection of relevant information without any additional cost and ultimately allowing the development of more robust biotechnological optimization efforts.

## 5. Conclusions

Photosynthesis is affected by the available irradiance and understanding how the former responds to the latter has both important natural and biotechnological implications.

Culture’s density has a substantial effect on microalgae photosynthetic activity. The highest the cell’s light absorption capacity, the greatest the effect on photosynthesis. Reducing the cell’s pigment content has been demonstrated to bring an advantage in dense microalgae cultures indeed, likely because of a more homogeneous light distribution profile, provided the environmental conditions drive to a relevant shading effect in WT cultures. High-sensitivity oxygen measurements are informative to understand how photosynthesis responds to culture density, opening the possibility to extrapolate already at the lab scale performances of industrial microalgae cultures and speeding up the identification of robust biological targets for biotechnological optimization.

## Acknowledgements

GP acknowledges the support from the University of Padova STARS Project WWBiomass. TM acknowledges the support from European Union H2020 Project No. 859770-NextGen-O2k and Marie Skłodowska-Curie No. 955520 Digitalgae.

