## Supplementary material for "Assessment of photosynthetic activity in dense microalgae cultures using oxygen production"

**Supplementary Tables**

**Supplementary Table S1. Estimation of photosynthetic parameters at different cell concentrations.** $\text{I}_{\text{comp}}$ and $\text{I}_{\text{sat}}$ indicate the light compensation and light saturation points, respectively, expressed as [μmol photons·s^-1^·m^-2^]. $\text{R}_{\text{D}}$ is the dark O₂ respiration rate, expressed as [pmol·s^-1^·10^-6^ cells], $\text{P}_{\text{gmax}}$ is the maximal gross O₂ photosynthetic rate, expressed as [pmol·s^-1^·10^-6^ cells]. $\text{Φ}_{\left( \text{I}_{\text{0}}\text{\_}\text{I}_{\text{comp}} \right)}$ is the quantum yield in the range between $\text{I}_{\text{0}}$ and$\text{ }\text{I}_{\text{comp}}$ [pmol O₂·m^2^·μmol photons^-1^·10^-6^ cells]. Data refer to the average ± SD of four independent biological replicates. Highlighted in yellow is the data from the two cell concentrations used in this work to represent a reliable description of diluted (5 · 10^6^ cells ml^-1^) and dense (100 · 10^6^ cells ml^-1^) microalgae cultures.

| **Cell concentration**  **(10^6^ cells mL^-1^)** | $\text{Φ}_{\left( \text{I}_{\text{0}}\text{\_}\text{I}_{\text{comp}} \right)}$ | **I_comp_** | **I_sat_** | **R_D_** | **P_gmax_** |
| --- | --- | --- | --- | --- | --- |
| **1** | 0.36 ± 0.10 | 14.6 ± 0.5 | 133 ± 59 | 5.28 ± 1.29 | 9.9 ± 1.2 |
| **2.5** | 0.09 ± 0.02 | 19.5 ± 2.3 | 153 ± 21 | 1.82 ± 0.21 | 4.9 ± 0.2 |
| **5** | 0.11 ± 0.01 | 12.2 ± 1.1 | 163 ± 27 | 1.31 ± 0.12 | 4.6 ± 0.2 |
| **10** | 0.1 ± 0.02 | 9.5 ± 1.2 | 149 ± 37 | 0.95 ± 0.16 | 4.3 ± 0.3 |
| **20** | 0.09 ± 0.01 | 9.9 ± 0.7 | 157 ± 26 | 0.85 ± 0.12 | 4.3 ± 0.2 |
| **40** | 0.09 ± 0.01 | 8.1 ± 1.0 | 175 ± 28 | 0.74 ± 0.17 | 4.0 ± 0.6 |
| **75** | 0.06 ± 0.01 | 10.7 ± 1.1 | 329 ± 288 | 0.62 ± 0.1 | 4.3 ± 1.2 |
| **100** | 0.05 ± 0.01 | 11.9 ± 1.1 | 597 ± 263 | 0.57 ± 0.09 | 4.3 ± 1.2 |

**Supplementary Table S2. Statistical analysis of correlation.** Values of the linear correlation equations for the datasets presented in panels c), d), e) and f) of Figure 5. Equations are expressed in the form: y = a + bx, with a, intercept and b, slope. R^2^ > 0.9 was used as threshold for linear relationships. Asterisk indicates when there is linear correlation between x and y observations, namely when the slope of the equations is statistically different from 0 (t-Test, p-value < 0.05).

| **Dataset (panel #)** | **Intercept** | **Slope** | **Pearson’s R** | **R^2^** |
| --- | --- | --- | --- | --- |
| **Fig. S4 - panel a** | -3.8 ± 26.4 | 0.2 ± 1.6 | 0.1 | 0.01 |
| **Fig. S4 - panel b** | 0 ± 0.1 | -0.55 ± 0.13 | -0.93 | 0.86 |
| **Fig. 5 - panel c** | 1.65 ± 7.1 | -0.25 ± 0.46 | -0.35 | 0.12 |
| **Fig. 5 - panel d *** | 0.12 ± 0.78 | -1.1 ± 0.05 | -0.99 | 0.98 |

**Supplementary Table S3. Estimation of photosynthetic parameters for all the strains used in this work at different cell concentrations.** $\text{I}_{\text{comp}}$ and $\text{I}_{\text{sat}}$ indicate the light compensation and light saturation points, respectively, expressed as [μmol photons·s^-1^·m^-2^]. $\text{R}_{\text{D}}$ is the dark O₂ respiration rate, expressed as [pmol·s^-1^·µg Chl], $\text{P}_{\text{gmax}}$ is the maximal gross O₂ photosynthetic rate, expressed as [pmol·s^-1^· µg Chl]. $\text{Φ}_{\left( \text{I}_{\text{0}}\text{\_}\text{I}_{\text{comp}} \right)}$ is the quantum yield in the range between $\text{I}_{\text{0}}$ and$\text{ }\text{I}_{\text{comp}}$ [pmol O₂·m^2^·μmol photons^-1^· µg Chl]. Data are normalized on the same Chl content and cannot be directly compared to those of Table S1. Traces used in this analysis are reported in Supplementary Figure S3. For each strain, alphabet letters indicate significant differences from diluted to dense conditions (Test-t, p-value<0.05). For the same culture condition, asterisks indicate significant differences between mutants and WT strain (t-Test, p-value<0.05). Data refer to the average ± SD of four independent biological replicates.

| **Strain ID** | **Cell concentration**  **(10^6^ cells mL^-1^)** | $\text{Φ}_{\left( \text{I}_{\text{0}}\text{\_}\text{I}_{\text{comp}} \right)}$ | **I_comp_** | **I_sat_** | **R_D_** | **P_gmax_** |
| --- | --- | --- | --- | --- | --- | --- |
| **WT** | **5** | 1.62 ± 0.18 | 14.2 ± 1 | 250 ± 29.5 | 23 ± 2.5 | 98 ± 3.3 |
| **WT** | **100** | 0.48 ± 0.05^a^ | 6.6 ± 0.8^b^ | 372 ± 71^c^ | 3.2 ± 0.5^d^ | 63 ± 2.8^e^ |
| **E2** | **5** | 1.46 ± 0.15 | 19.1 ± 1.1* | 256 ± 27.7 | 28 ± 2.9* | 104 ± 3.8 |
| **E2** | **100** | 0.57 ± 0.05^a^ | 6.6 ± 0.8^b^ | 364 ± 67.6^c^ | 3.8 ± 0.6^d^ | 72 ± 3.2*^e^ |
| **I2** | **5** | 1.8 ± 0.2 | 15.8 ± 1.1* | 243 ± 25 | 29.5 ± 3* | 113 ± 3.8* |
| **I2** | **100** | 0.51 ± 0.05^a^ | 8.6 ± 0.5*^b^ | 493 ± 220^c^ | 4.4 ± 0.6*^d^ | 77 ± 8.4*^e^ |
| **I48** | **5** | 1.3 ± 0.08 | 27.3 ± 0.6* | 330 ± 38* | 34 ± 1.9* | 114 ± 2.9* |
| **I48** | **100** | 0.5 ± 0.04^a^ | 7 ± 0.7^b^ | 831 ± 38*^c^ | 3.4 ± 0.5^d^ | 86 ± 25.8 |
| **I29** | **5** | 1.7 ± 0.2 | 17 ± 1.2* | 237 ± 22 | 28.7 ± 3* | 107 ± 3.8* |
| **I29** | **100** | 0.6 ± 0.06^a^ | 8.5 ± 0.6*^b^ | 406 ± 101^c^ | 5 ± 0.7*^d^ | 86 ± 5^e^* |

**Supplementary Figures**

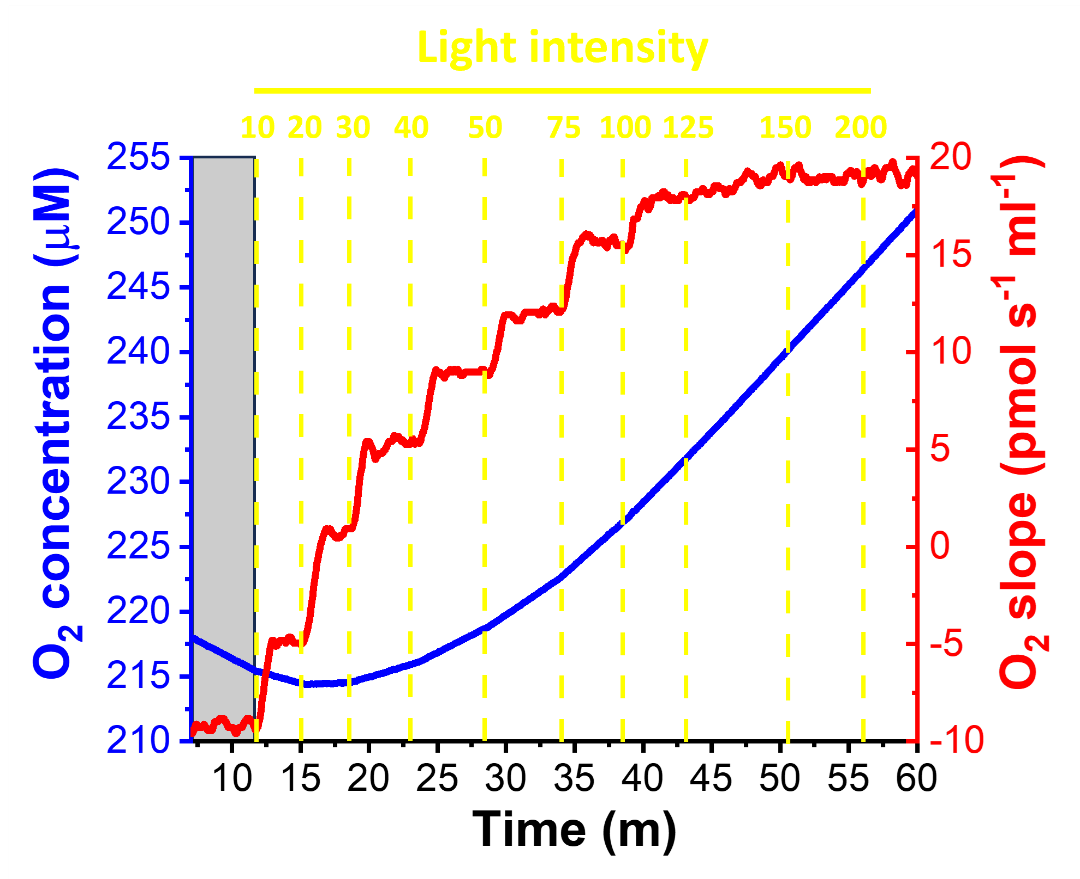

**Figure S1. Raw data collected for building a PI relationship.** Raw data obtained from a typical experiment with the high-sensitivity Clark electrode employed in this work. When the light intensity increases (vertical dashed yellow lines, values are expressed as μmoles of photons m^-2^ s^-1^), the O_2_ slope (i.e. time-derivative of oxygen concentration in the sample) first increases then stabilizes. 40 data points were collected when the O_2_ slope trace was stable are averaged and used to build the plot of Figure 1. The grey box indicates the initial dark respiration phase.

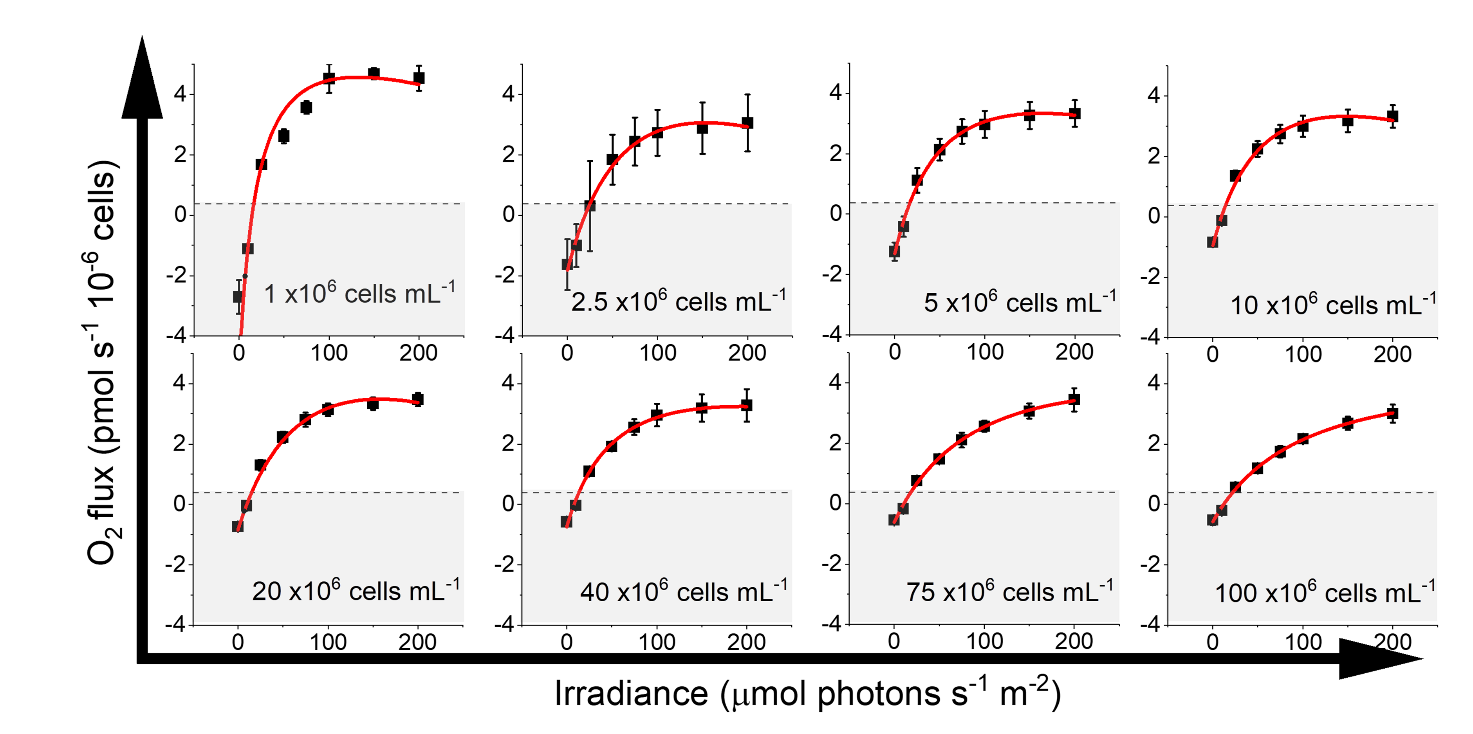

**Figure S2. Photosynthesis-Irradiance relationship as a function of cell concentration.** Photosynthesis is expressed as O_2_ flux and it is normalized to the number of cells. Data were fitted with the equation defined in (Ye, 2007) using a minimum mean square error-based approach (red curves). In each panel, the area where the respiration rate is higher than photosynthesis is marked in grey. The photosynthetic parameters obtained from each curve are detailed in Supplementary Table S1. Data refer to the average ± SD of four independent biological replicates.

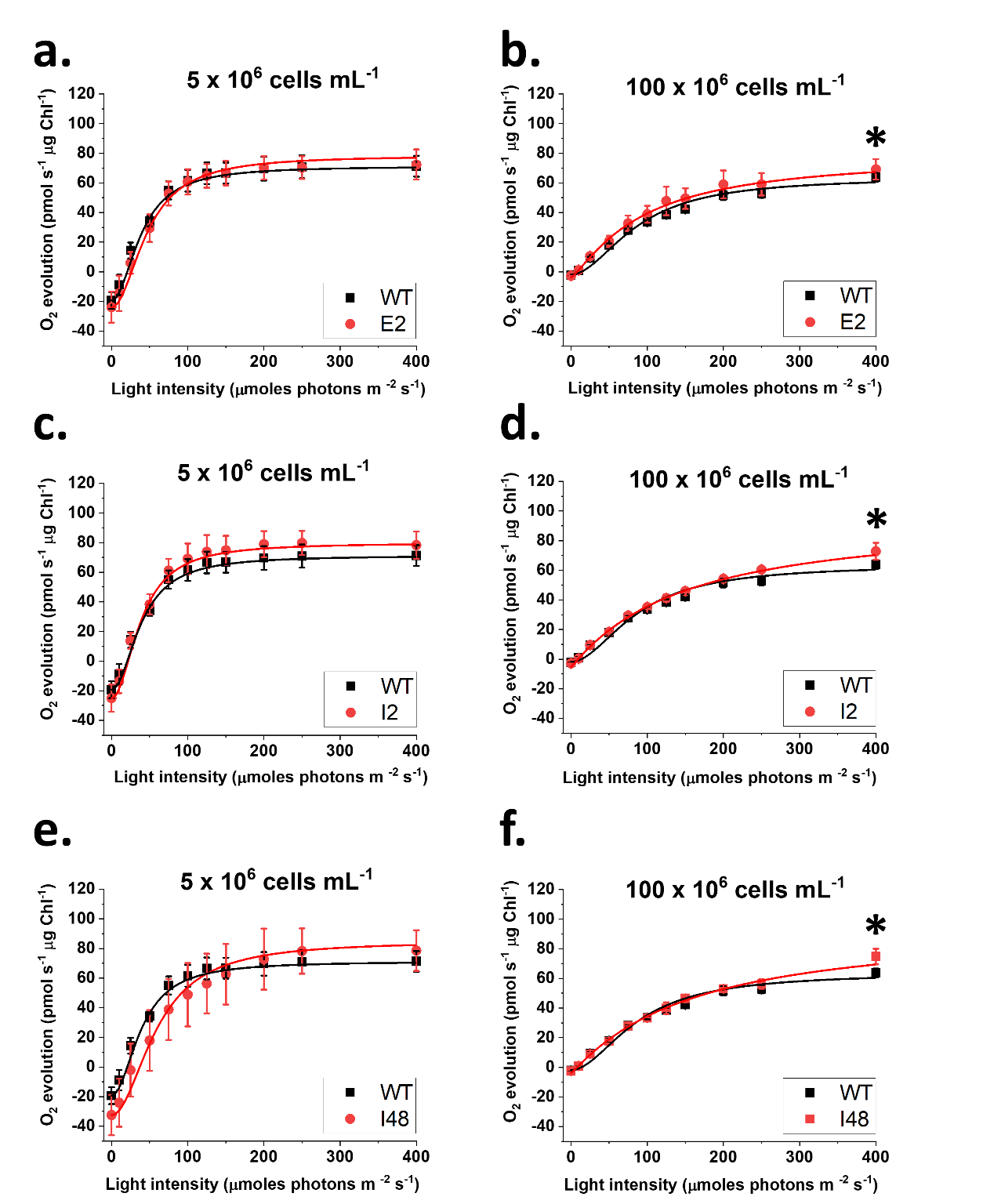

**Figure S3. Photosynthesis-Irradiance relationship in pale green *Nannochloropsis* mutants.** Photosynthesis is expressed as O_2_ flux and it is normalized to the same pigment content to account for the different absorption properties of the different strains. Data were fitted with the equation defined in (Ye, 2007) using a minimum mean square error-based approach (red and black curves). Data refer to the average ± SD of four independent biological replicates. Asterisks indicate statistically significant differences between one mutant and the parental strain (t-Test, p-value < 0.05). A, c and e show the PI curves of strain E2, I2 and I48 (in red), respectively, compared to the WT (in black) in diluted cultures (i.e. 5 · 10^6^ cells ml^-1^). B, d and f show the PI curves of strain E2, I2 and I48 (in red), respectively, compared to the WT (in black) in dense cultures (i.e. 100 · 10^6^ cells ml^-1^).

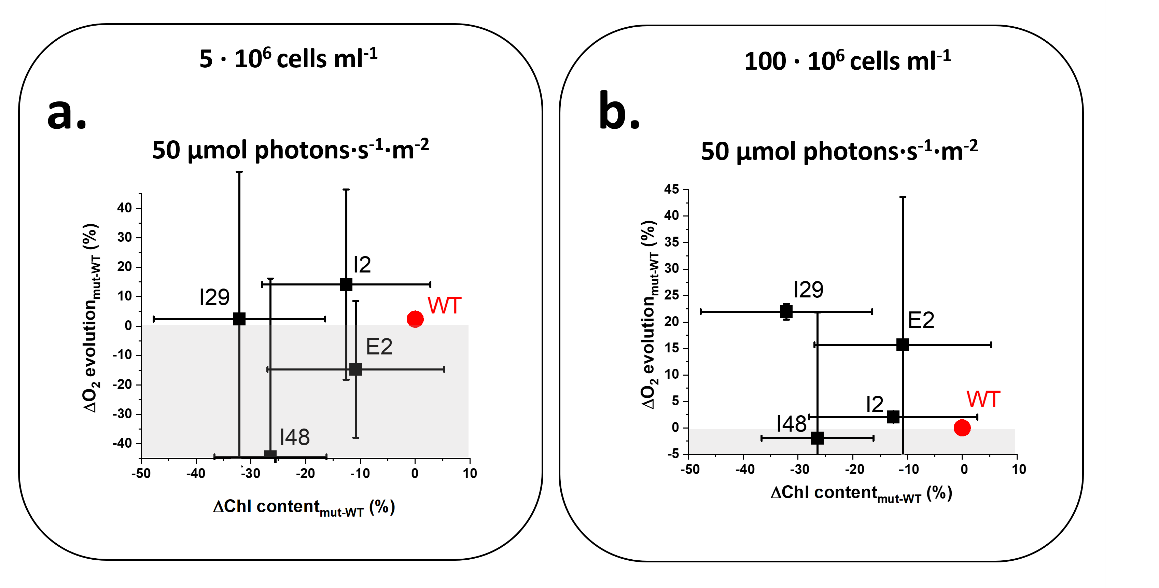

**Figure S4. Improvement of photosynthetic activity as a function of the reduction of the pigment content, in *Nannochloropsis* cultures with different densities.** Improvement in O_2_ flux at 50 μmol photons s^-1^ m^-2^, expressed as percentage with respect to the value of the parental strain (WT) for the four mutants showing different degrees of reduction in the chlorophyll (Chl) content, expressed as percentage of reduction with respect to the WT, for diluted (a) and dense (b) *Nannochloropsis* cultures. Data correspond to the photosynthetic activity measured at 50 μmol photons·s^-1^·m^-2^, that is the light intensity at which there is linear correlation between photosynthetic activity and irradiance (Figure 1). Grey areas refer to a lower photosynthetic activity than the WT. There isn’t linear correlation between x and y observations in both panels (t-Test, p-value < 0.05). Parametric values for the correlation equations of both panels are reported in Supplementary Table S2. Data refer to the average ± SD of four independent biological replicates. The original data used in this figure are reported in supplementary figure S3.
